# Machine learning to classify mutational hotspots from molecular dynamic simulations

**DOI:** 10.1101/2023.09.07.556625

**Authors:** James Davies, Georgina Menzies

## Abstract

Benzo[*a*]pyrene, a notorious DNA-damaging carcinogen, belongs to the family of polycyclic aromatic hydrocarbons commonly found in tobacco smoke. Surprisingly, nucleotide excision repair (NER) machinery exhibits inefficiency in recognising specific bulky DNA adducts including Benzo[*a*]pyrene Diol-Epoxide (BPDE), a Benzo[*a*]pyrene metabolite. While sequence context is emerging as the leading factor linking the inadequate NER response to BPDE adducts, the precise structural attributes governing these disparities remain inadequately understood. We therefore combined the domains of molecular dynamics and machine learning to conduct a comprehensive assessment of helical distortion caused by BPDE-Guanine adducts in multiple gene contexts. Specifically, we implemented a dual approach involving a random forest classification-based analysis and subsequent feature selection to identify precise topological features that may distinguish adduct sites of variable repair capacity. Our models were trained using helical data extracted from duplexes representing both BPDE hotspot and non-hotspot sites within the *TP53* gene, then applied to sites within *TP53, cII*, and *lacZ* genes.

We show our optimised model consistently achieved exceptional performance, with accuracy, precision, and f1 scores exceeding 91%. Our feature selection approach uncovered that discernible variance in regional base pair rotation played a pivotal role in informing the decisions of our model. Notably, these disparities were highly conserved among *TP53* and *lacZ* duplexes and appeared to be influenced by the regional GC content. As such, our findings suggest that there are indeed conserved topological features distinguishing hotspots and non-hotpot sites, highlighting regional GC content as a potential biomarker for mutation.

**Author Summary:** Although much is known about DNA repair processes, we are still lacking some fundamental understanding relating to DNA sequence and mutation rates, specifically why some sequences mutate at a higher rate or are repaired less than others. We believe that by using a combination of Molecular Simulation and Machine Learning (ML) we can measure which structural features are present in sequences which mutate at higher rates in cancer gene and lab-based test assays frequently used to investigate toxicology.

Here we have run Molecular Dynamics on five sets of DNA sequences with and without a carcinogen found in cigarette smoke to allow us to study the mutation event that would need to be repaired. We have measured their helical and base stacking properties. We have used ML to successfully differentiate between low and high mutating sequences using this model allowing us to begin to elucidate the structural features these groups have in common.

We believe this method could have wide reaching uses, it could be applied to any gene context and mutation event and indeed the knowledge of the structural features which are best repaired gives us insight into the biophysics of DNA repair adding knowledge to the drug design pipeline.

## Introduction

Mutations serve as the fundamental source of genetic variation, fostering adaptive evolution, whilst simultaneously contributing to the development of diseases such as cancer and age-related illnesses [1,2]. Though random, a nucleotides mutation rate, defined by its probability to undergo mutation, exhibits considerable heterogeneity throughout the genome. This variation has been demonstrated at various levels, with the rate of site-to-site mutation shown to vary by more than 100-fold across the genome, prompting sequence dependant contribution to both evolution and genetic pathologies [3]. Analysing the precise features that promote this dependency, as such, may unlock valuable insights into the mutagenic mechanisms underlying various human diseases.

It is widely accepted that site-specific mutational frequencies are contingent upon three generalised factors: (i) nucleotide stability and their vulnerability to mutagenesis; (ii) the fidelity of DNA replication processes; and (iii) the efficiency of DNA repair machinery [4]. Consequently, a natural gradient manifests, revealing the degree to which mutational occurrences are influenced by the underlying sequence composition. For instance, regions of repetitive DNA such as homonucleotide runs and microsatellite repeats serve as an active source of hypermutability through polymerase slippage events, emphasising motif recurrence over precise sequence context [5,6]. In contrast, alternative mutation spectra exhibit pronounced sequence specificity, often guided by intricate local and occasionally distant sequence contexts [7-11]. While the underlying causes of the latter remain subjects of substantial debate, emerging evidence indicates that this interdependence forms the foundation for numerous disease-associated mutation spectra. This is notably evident in the case of the *TP53* mutation spectrum within the context of lung cancer [12,13].

*TP53* is a gene that is responsible for encoding the p53 tumour suppressor protein, commonly referred to as the “Guardian of the Genome” [14]. Its principal biological role involves safeguarding DNA integrity through the inhibition of cell proliferation and the promotion of apoptosis in response to cellular stress [15]. This renders *TP53* a pivotal target for inactivation in diseases, consequently establishing it as the most commonly mutated gene in human cancer [12]. *TP53* mutations are particularly prevalent in lung cancer, with approximately 50% of tumours containing a mutation [16]. Mutation distribution, however, is shown to vary substantially between lung and most other tumour types, with a higher frequency of G:C>T:A transversions in the former [12,16,17]. While the influence of sequence context on the varying *TP53* mutation spectrum continues to be a subject of intense debate, a consensus exists regarding the underlying cause: the formation of DNA adducts induced by polycyclic aromatic hydrocarbons (PAH) found in cigarette smoke [18-20].

Benzo[a]pyrene (B[*a*]P) emerges as the predominant PAH stemming from the incomplete combustion of organic material within tobacco smoke. A study by Vu *et al*., established that one cigarette can yield an intake of 13-38ng of B[*a*]P [21]. This compound is then metabolically processed to generate a series of carcinogenic derivatives, notably including the trans(+)anti-benzo[a]pyrene diol epoxide, hereon referred to as BPDE (Figure 1A) [22]. Empirical evidence suggests that a combination of preferential DNA damage and poor repair is responsible for the lung associated mutation hotspots in *TP53* [23]. Recurrently mutated codons 157, 158, 245, 248, 273, and 282 harbour methylated CG dinucleotides and therefore represent favourable locations for BPDE binding while concurrently exhibiting resistance to nucleotide excision repair (NER) mechanisms [24]. Notably, the (K)-7S,8R,9R,10S□+=□anti-B[*a*]PDE enantiomer (10S) DNA adduct dominates, accounting for an overwhelming majority (70.8-92.9%) of the total adducts at these specific sites (Figure 1B-C) [25]. Interestingly, however, Denissenko *et al*., report other known targets of BPDE, including 170, 186, 202, 213, 267 and 290, whose mutation is either uncommon or silent in lung cancer [23,26,27]. Given the broad specificity of BPDE and accounting for selective processes, the induction of *TP53* mutations by BPDE likely the combined result of differential adduct availability and varying repair capabilities at distinct adduct sites.

**Figure 1:**
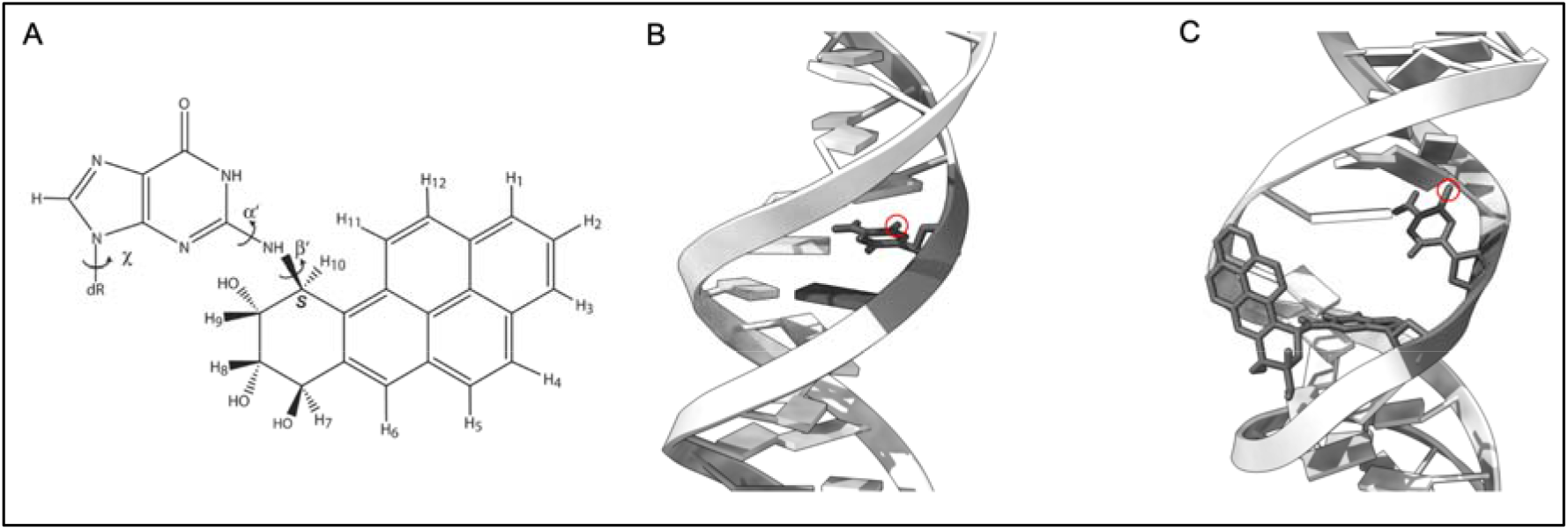
Chemical structure and mutagenic adduct formation of Benzo[*a*]pyrene diolepoxide. (**A**) Chemical structure of 7S,8R,dihydroxy-9R,10S-epoxy-7S,8R,9R,10S (+)-trans-anti-B[*a*]PDE; (**B**) regular unadducted DNA with methyl group (highlighted by a red circle) on the cytosine at base 6; (**C**) adduct of BPDE (sticks) bound to guanine in a CpG site in minor groove directed towards the 5′ end of the DNA sequence.

Importantly, while NER is capable of repairing a diverse range of bulky lesions, its efficiency varies dramatically according to the chemical structure, stereochemistry, confirmation, and sequence context surrounding the lesion site [28-32]. Various studies, both experimental and theoretical, highlight how variable sequence context can influence NER efficacy by impacting DNA structure and subsequent recognition processes [29,32-34]. Paul *et al*., reveal that distinct resistances to NER-induced untwisting/bending renders certain contexts more likely to access conformations required for successful repair, thereby proposing a sequence-dependent propensity and trajectory for DNA “unwinding” [35]. Furthermore, we previously reported sequence-dependent variability in DNA distortion at BPDE adduct sites linked to lung cancer-associated *TP53* mutation sites [36]. Thus, the potential disparity in repair capacity within *TP53* across BPDE adduct sites may arise from sequence-driven modifications in DNA structural parameters. Despite these notable strides, however, the true extent to which sequence context influences the distortional profiles induced by BPDE adducts, and consequently, NER, remains a subject of ongoing debate, prompting the need for continued scientific scrutiny.

In this study, we sought to integrate the realms of molecular dynamics (MD) and machine learning (ML) to formulate a model capable of defining the precise features that promote the differing mutational pattern observed in *TP53* during lung tumorigenesis. We hypothesised that by the inclusion of ML methodologies we would significantly amplify the range of hotspots sites under analysis, thereby enabling the comparison of mutation spectra within single or multiple gene contexts. Our initial aim was to determine whether helical information acquired from MD simulation could indeed be used to isolate sequence dependant features that aberrate the NER of BPDE lesions in *TP53*. Upon achieving this, our subsequent goal was to ascertain the model’s efficacy in utilising this information in predicting the mutation status of unseen helical data from alternative gene contexts that are recognised to undergo BPDE-induced mutations including *cII* and *lacZ*. The knowledge gained could help provide more information surrounding the aetiology of the disease, and allow for the develop innovative anti-cancer therapies aimed at reinstating healthy NER function at these sites.

## Results

### 2.1 Data Retrieval and Definition of Spectral Investigation

Our analysis was focused on mutational spectra of three distinct gene contexts – *TP53, cII*, and *lacZ* – each exhibiting variable repair of BPDE-induced lesions as defined by [37], [38], and [39]. Mutational data obtained for *TP53* was used to assemble three further datasets. The datasets, named “TP53”, and a secondary dataset obtained from unique simulations of corresponding *TP53* sequences, termed “Val,” were constructed based on *TP53* duplexes with site-restricted methylation, serving as the training and validation sets, respectively, for our ML model. Subsequently, a third dataset named “Meth” was compiled to assess the impact of more distal methylation on *TP53* dynamics, through homomethylation. Mutational profiles derived from the reporter genes *cII* and *lacZ* served as the foundation for subsequent investigations, allowing us to infer which components (if any) appear to have a conserved role or association with sample characteristics, such as repair phenotype or adduct binding state.

### 2.2 Conformational Stability and Sequency

Root mean square deviation (RMSD) and total energy analysis provided confidence about the stability of the simulations. Elevated RMSD values typically suggest that flexibility and movement during a simulation are likely characterised by instability. For *TP53*, total energy calculations confirmed the stability of all simulations conducted (Figure 3A). Average RMSD values for each sequence, relative to the initial structure, provided valuable insights into the variations in flexibility observed throughout the 100ns simulation period. As depicted in Figure 3B, unbound *TP53* duplexes showed a mean RMSD of 3.83Å ± 0.81, while adducted sequences exhibited a higher mean RMSD of 5.50Å ± 1.10. Fluctuations around the mean remained stable throughout the simulation period of 10-100ns, which prompted the exclusion of the initial 10ns from all subsequent analyses. Additionally, the results obtained from the RMSD analysis demonstrated a consistent trend of increased flexibility in the adducted sequences compared to the non-adducted controls. This observation aligns with our expectations, as BPDE adducts are known to disrupt the helical structure, resulting in increased fluctuation and mobility. All differences in the distributions of RMSD values between adducted and control sequences were significant (*P* < 0.000).

**Figure 2:**
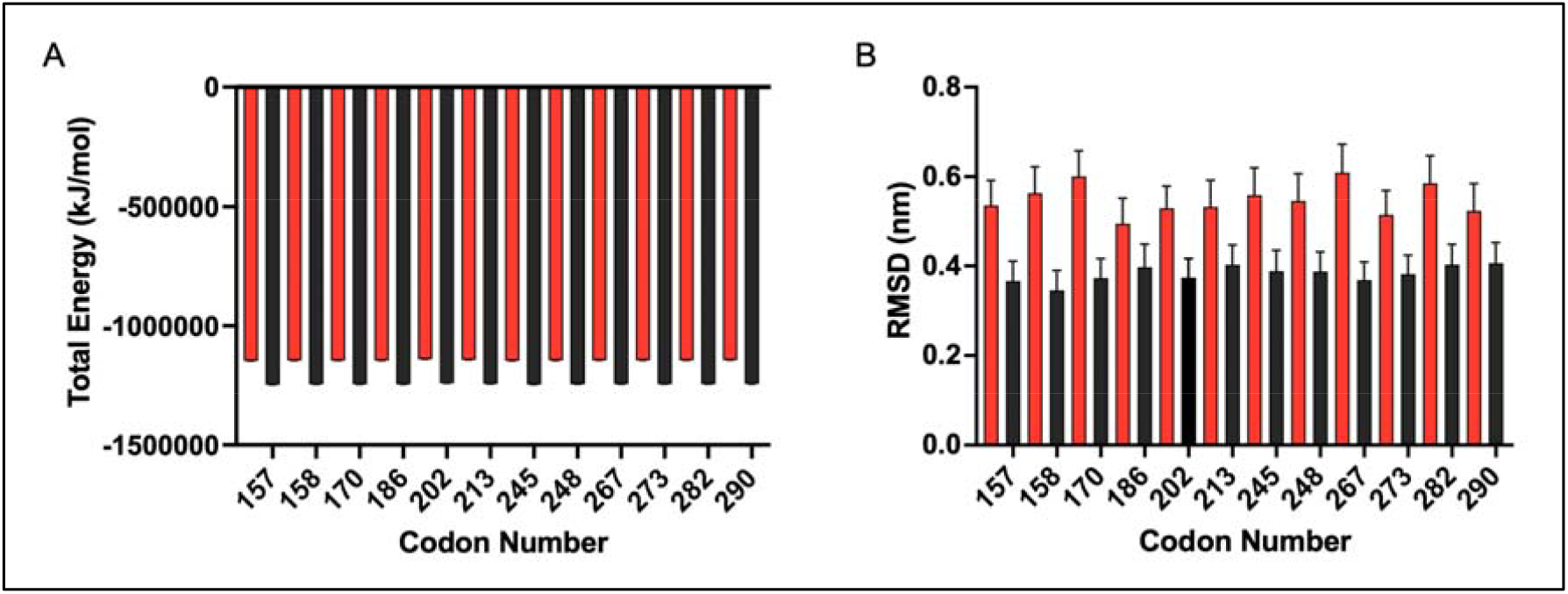
Conformational stability and flexibility of adducted and control TP53 sequences. Total energy (**A**) in KJ/mol and RMSD values (**B**) in nm for control sequences (black) and adducted sequences (red); error bars show ± standard deviation. Corresponding values for Val, Meth, cII, and lacZ datasets are depicted in Supplementary Figures S1-S4.

**Figure 3:**
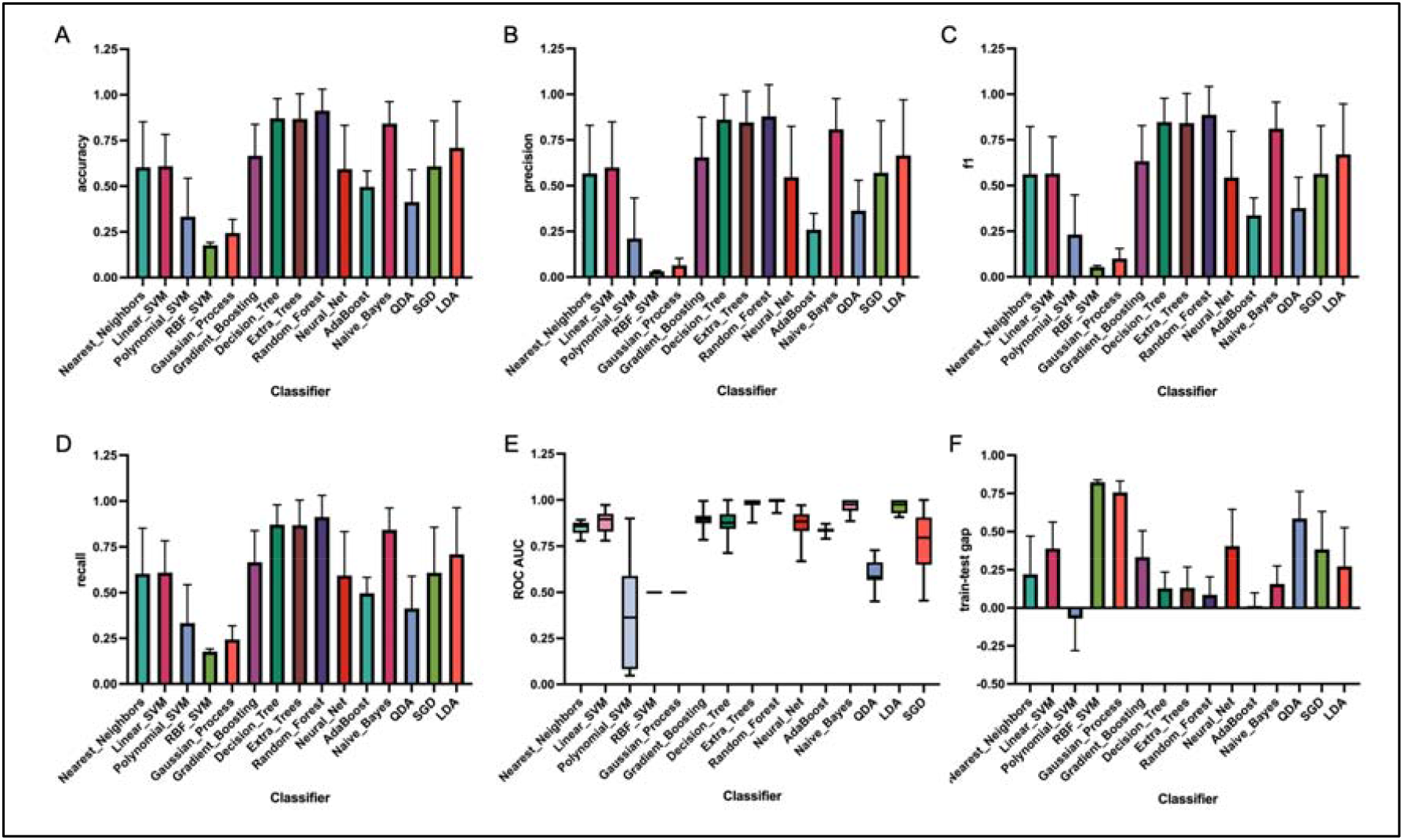
Evaluation results of fifteen classification algorithms using accuracy (A), precision (B), f1 (C), recall (D), ROC AUC, and training-test gap scores (F). Error bars represent ± standard deviation for panels (A) to (D) and indicate the range of ROC AUC scores in panel (E), with the median score depicted by a solid black line.

### 2.3 Classifier Comparison

To select an appropriate classifier, we first compared the prediction results of 15 unique classification algorithms on the TP53 dataset. Each classifier was instantiated with default hyperparameters, and model performance was evaluated through the quantification of accuracy, precision, recall, f1, and area under the curve (AUC) and receiver operating characteristic (ROC) scores. Among the 15 algorithms, decision tree-based models (Decision Tree, Extra Trees, and Random forest (RF)) were by far the most prominent, demonstrating advanced results ranging from 0.84 to 0.91 across each evaluation metric (Figure 4A-D). Kernel machines (Polynomial (support vector machine) SVM, radial basis function (RBF) SVM, and Gaussian Process), conversely, encountered substantial challenges during the classification process, resulting in a notable decline in accuracy, precision, recall, f1, and, in the case of Polynomial SVM, considerable variation in ROC AUC (Figure 4E). As shown in Figure 4A-F, the RF algorithm exhibited the best discriminatory power, with all mean evaluation scores within the range of 0.88-0.99. ROC AUC values displayed minimal range (±0.035), highlighting the models consistent and robust discrimination across different scenarios (Figure 4E). We observed that, in comparison to all other well-performing algorithms, the RF model exhibits the smallest training-test gap (Figure 4F). This finding confirms the superior generalisation capacity of the RF model, solidifying its standing as the optimal classifier for our study.

**Figure 4:**
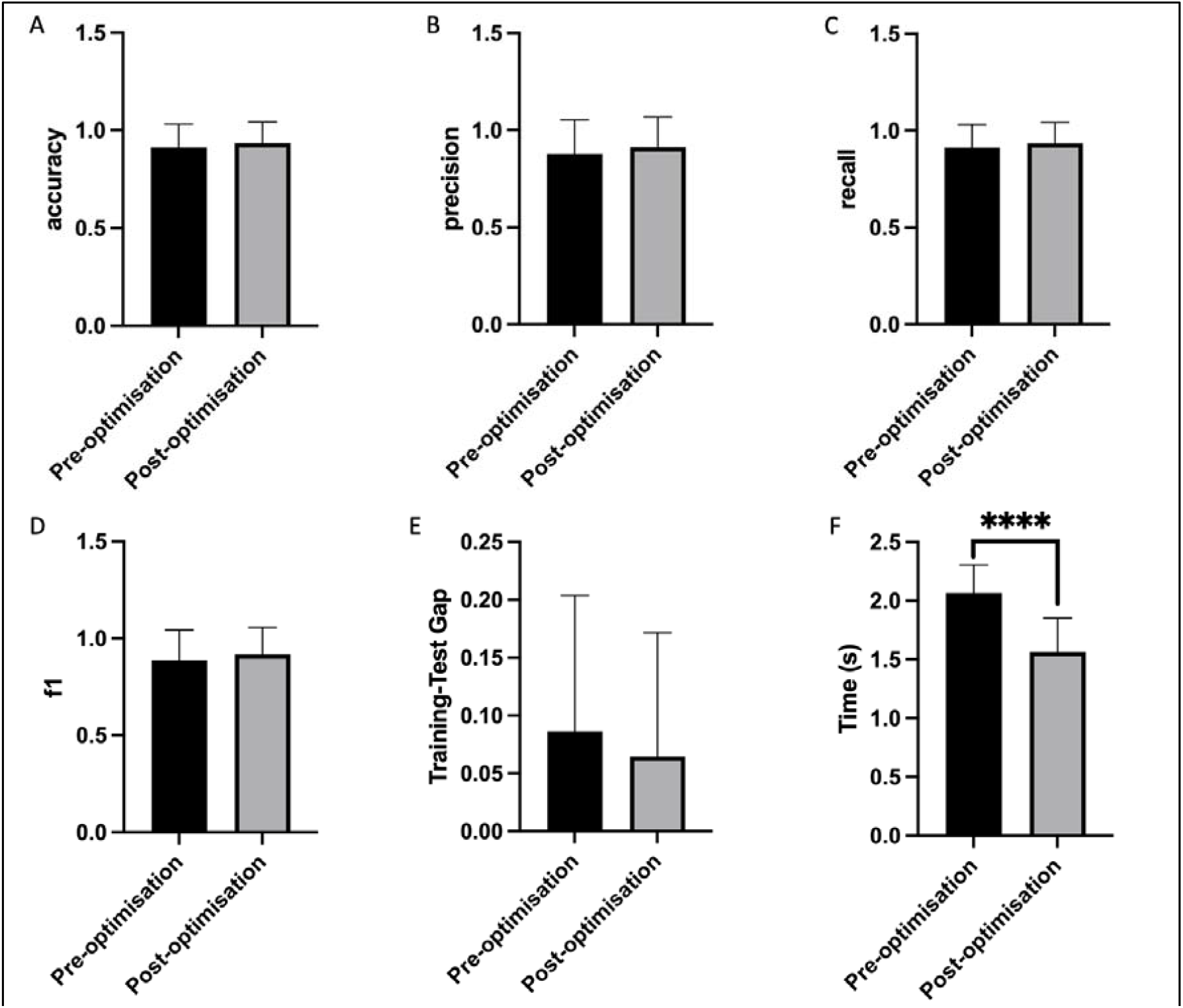
RF optimisation by GSO. Bar chart Illustrating the averaged accuracy (A), precision (B), recall (C), f1 (D), train-test gap (E), and execution time (F) scores before and after optimisation. Pre-optimisation scores are shown in black, while post-optimisation scores are depicted in grey. Error bars indicate ± standard deviation.

### 2.4 Grid Search Optimisation

To identify the optimal hyperparameter configuration of our model, we employed Grid Search Optimization (GSO). Crucial parameters, including bootstrap, min_sample_leaf, max_features, and n_estimators, significantly influence the performance of the RF classifier. Utilising GSO, we transformed these parameters into a grid with a defined spatial range and explored all points to obtain the global optimal solution.

Incorporating novel GSO parameters into the existing RF algorithm was shown to enhance model performance, yielding a 3.06% increase in model accuracy and 12.63% improvement in precision (Figure 4A,B). Hyperparameter tuning also induced modest improvements in recall, f1, and training-test gap scores (Figure 4C-E). This not only bolstered overall accuracy but also honed the model’s ability to detect positive cases while mitigating instances of false negatives and false positives. Further to this, GSO was shown to significantly improve the model’s operating speed (P<0.0001) (Figure 4F). This is to be expected given that GSO typically accelerates convergence during model training.

### 2.5 *TP53* Mutation

To elucidate the potential relationship between structural distortion and mutation frequency within the context of *TP53*, we initially deployed the optimised RF model. This approach evaluated the model’s ability to leverage the knowledge acquired during the original training phase to classify two previously unseen datasets. One dataset, sourced from the re-simulation of previously utilised *TP53* starting structures, constituted the designated validation dataset. Additionally, we simulated novel starting structures representative of the same *TP53* sequences, characterised, however, by full CpG methylation, thereby creating the methylated dataset. Once more, model performance was scrutinised using evaluation metrics accuracy, precision, f1 and recall, the values of which are depicted in Table 1. Feature scoring was then applied exclusively to adduct-bound sequences to deduce the predictive utility of each helical parameter for sample classification.

**Table 1:**
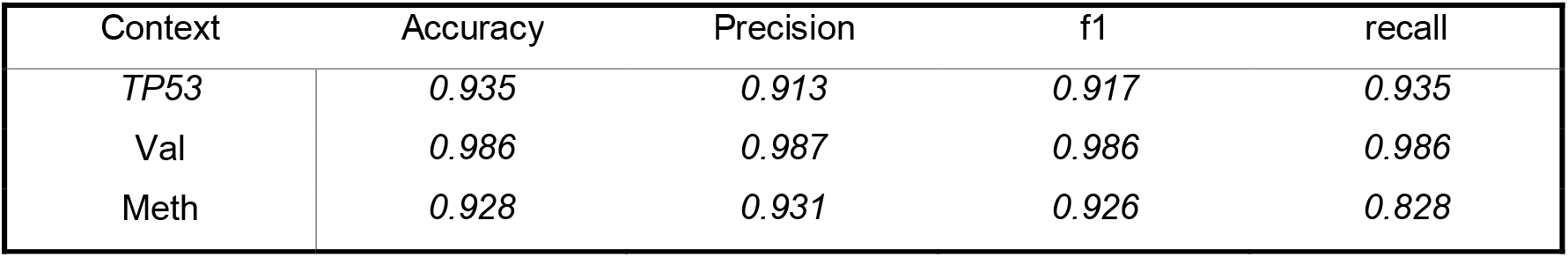
Mean accuracy, precision, f1 and recall scores for RF model using 10-fold cross-validation.

| Context | Accuracy | Precision | f1 | recall |
| --- | --- | --- | --- | --- |
| *TP53* | 0.935 | 0.913 | 0.917 | 0.935 |
| Val | 0.986 | 0.987 | 0.986 | 0.986 |
| Meth | 0.928 | 0.931 | 0.926 | 0.828 |

Table 1 reveals a prominent improvement in our model’s performance upon its deployment with validation data. The model achieves a near perfect performance, showcasing an average recall, accuracy, and f1 score of 0.986, coupled with an average precision score of 0.987. Notably, precision and f1 metrics exhibit an approximate 8% increase, indicating improved accuracy in positive predictions and a refined equilibrium between precision and recall. Despite the introduction of added complexity in the form of CpG methylation, our RF model maintains exceptional classification performance, achieving high scores in precision (0.931), f1 (0.926), and accuracy (0.928). Prediction recall scores, however, experience a marginal decline, registering a modest value of 0.828, indicating potential challenges in accurately identifying positive instances in the event of differentially methylated training and test datasets.

Feature scoring unveiled notable alignments in parameter types defined as pivotal during the classification process. Predominantly, parameters illustrating the rotation of base pairs/base pair axes took precedence among the top 10% of scored features, contributing to more than 61% of highlighted components (Table 2). Notably, a distinctive bias was unveiled towards the axis-displacement parameter tip, constituting 25% and 19% of the identified features in the *TP53* and validation datasets, respectively. Inclination and tip angles intricately mirror the orientation of base-pair planes concerning the helical axis. This observation naturally prompts the anticipation that parameters governing base step planarity such as roll, tilt, and buckle angles would also exhibit elevated scoring. Our dataset validates this expectation by showcasing a substantial prevalence of these parameters within the upper echelons of the top 10% of featured attributes.

**Table 2:** Top 10 helical parameters within TP53-related datasets identified by feature scoring. Abbreviations explained in supplementary table four.

| Rank | *TP53* | Val | Meth |
| --- | --- | --- | --- |
| 1 | 14pl | 23re | 14pl |
| 2 | 23re | 14pl | 21ad |
| 3 | 10ad | 23tp | 13pl |
| 4 | 10tp | 13pl | 14og |
| 5 | 11st | 11st | 11tp |
| 6 | 10xp | 16ad | 23tp |
| 7 | 14sh | 10tp | 6pl |
| 8 | 13sg | 23sr | 12se |
| 9 | 17ad | 11pl | 13ad |
| 10 | 23tp | 10xp | 13se |

Encouragingly, these trends extended to the methylation dataset, with tip, buckle, axis-bend, propel, and opening continuing to dominate as primary helical attributes. Notably this set of five parameters accounted for an impressive 72% of identified features, with tip emerging as the most influential contributor, commanding 19% of the overall proportion. Bases 10, 14, and 23 maintained their position as the highest-scoring base positions across all three *TP53* datasets. However, due to their distal location from the lesion site, it remains challenging to draw definitive conclusions regarding the precise significance of these positions in relation to the observed distortion.

### 2.6 Alternative Gene Contexts

After deploying our RF model and feature scoring pipeline, we employed the approach to deduce the relative discriminatory potential of our training data concerning hotspot and non-hotspot samples in alternative gene contexts, and identify potentially conserved helical parameters likely to contribute to the repair phenotype. Analogous to the previous mutational analysis, we harnessed the insights gained from the initial training event to guide the investigation of two supplementary unseen gene contexts with established BPDE-binding sites—namely, *lacZ* and *cII*. Mean accuracy, precision, f1 and recall scores obtained from 10-fold cross-validation can be seen in Table 3.

**Table 3:** Mean accuracy, precision, f1 and recall scores for RF model using 10-fold cross-validation.

| Context | Accuracy | Precision | f1 | recall |
| --- | --- | --- | --- | --- |
| *cII* | 0.628 | 0.737 | 0.639 | 0.628 |
| *lacZ* | 0.788 | 0.884 | 0.800 | 0.788 |

As anticipated, deploying the RF model on structural data acquired from duplexes with unseen base compositions yielded a noticeable decline in classifier performance. Specifically, in the context of *cII*, accuracy, f1, and recall demonstrated substantial reductions of approximately 43-45%. Remarkably, despite these changes, precision scores remained steadfast at 0.737, highlighting the model’s classification potential amid shifting sequence contexts. Similarly, the RF model’s predictive capacity encountered limitations when applied to the *lacZ* dataset. The model accurately predicted a significant proportion of samples (78.8%), achieving a notable precision rate of 88.4%. Nonetheless, it’s worth noting that recall (78.8%) slightly lagged precision, hinting at potential cases where positive instances might not have been precisely identified. These observations align with our expectations, underscoring the model’s sensitivity to sequence intricacies and elucidate the potential boundaries surrounding the applicability of ML algorithms trained on individual gene contexts.

Examining the top-ranked features in *cII* and *lacZ* datasets reaffirmed a notable emphasis on rotation-specific parameters. Parallel to that observed in *TP53*, the tip parameter remained dominant, constituting 22% and 28% of identified features in *cII* and *lacZ* datasets, respectively (Table 4). Similarly, an abundance of parameters associated with base step rotation, including buckle, inclin, propel, and opening, persisted, accounting for 58% (*cII*) and 69% (*lacZ*) of the top 36 features in each dataset. Despite minor alterations in the distribution of rotational parameters, these findings closely resembled those within *TP53*-specific contexts, accentuating the emphasis on rotational parameters as primary contributors to the discrimination process. Additionally, it’s worth noting that the positioning of these parameters exhibited notable deviation from *TP53* results, aligning with our expectations due to the change in base contexts. Collectively, these insights underscore the fundamental role of rotational parameters in shaping the model’s discriminative prowess across diverse gene contexts.

**Table 4:** Top 10 helical parameters within non-TP53 datasets identified by feature scoring. Abbreviations detailed in supplementary table four.

| *Rank* | *cII* | *lacZ* |
| --- | --- | --- |
| 1 | 17rl | 10in |
| 2 | 15in | 9be |
| 3 | 13tp | 17tp |
| 4 | 16ad | 20sh |
| 5 | 8se | 16tp |
| 6 | 15ad | 22tt |
| 7 | 15tp | 10be |
| 8 | 11yp | 17be |
| 9 | 21tw | 17sg |
| 10 | 8in | 5tp |

## Discussion

The intricacies of DNA structure and its dynamic flexibility constitute essential features of sustained eukaryotic life [40-43]. Paradoxically, the very attributes that underscore DNA’s functionality can also support detrimental processes, such as those driving tumorigenesis [36,44,45]. While prior research has exposed the influence of BPDE on damaged base conformation and DNA bending, a thorough investigation into the structural factors driving mutations across multiple gene contexts has not been pursued [8,36,46]. To this end, we conducted a focused investigation to uncover how sequence context influences the structural modifications induced by BPDE adducts at mutation sites within *TP53, cII*, and *lacZ* genes. This allowed us to identify conserved patterns in helical distortion among regions of elevated mutational frequency, irrespective of gene context, offering novel insights that can aid in the development of anti-cancer therapies.

Accessing and interpreting the information embedded within MD data, is non-trivial due to its high volume and dimensionality. As such, we embraced an approach based on ML principals, facilitating a comprehensive appraisal of helical structure across a substantial number of duplex samples. With various ML methodologies offering unique advantages for analysing such datasets, we embarked on an evaluation of 15 distinct ML classifiers, aiming to pinpoint the most appropriate model for our specific framework. As anticipated, decision tree-based models, including Decision Tree, Extra Trees, and RF, consistently outperformed other models across different class variables. Although the performance differences among these three models was marginal, it is noteworthy that the RF algorithm consistently achieved the highest scores for all evaluation metrics, and so became our method of choice for further analysis.

We leveraged a well-established feature in the field of cancer biology – the *TP53* mutation spectrum – to evaluate our model’s proficiency in classifying previously unseen helical data [37]. When applied to the validation dataset, the model achieved the highest prediction scores, instilling confidence in the robustness and accuracy of our RF. Despite maintaining exceptional performance on the validation data, however, the classifier exhibited partial regression when applied to the methylation dataset. This variance is likely attributed to the differential methylation statuses between the training and test data, underscoring minor limitations in model generalisation. Indeed, the influence of CpG methylation on DNA flexibility and its potential repercussions on NER have been a subject of persistent inquiry. Several studies have indicated that regions with heightened cytosine methylation enhance NER function at BPDE lesions [47-49]. Given our models adaptability to methylation data, we suggest that cytosine methylation has limited influence on the observed dynamics, and therefore, on NER efficiency. This aligns seamlessly with the notion that all *TP53* CpG sites within lung tissue exhibit methylation, implying that methylation, in isolation, cannot comprehensively explain the variable DNA repair rates witnessed at BPDE binding sites [50].

The model’s effectiveness noticeably declined when confronted with helical data from different gene contexts, aligning with the notion that underscores model sensitivity to variations in contextual information. While it is commonly agreed that imbalanced datasets adversely impacts classifier performance, it is possible to mitigate population bias and improve overall model training by combining all available data [51]. However, this strategy can complicate the assessment of information specific to the training dataset. Thus, we made the deliberate choice to maintain the segregation of data. It’s essential to note that our input data had some limitations, particularly in the absence of information regarding non-hotspot sites. To address this, for *lacZ* we defined non-hotspot sites based on reported codon mutational frequencies upon BPDE exposure, categorising them as sites with lower mutational frequencies compared to those of defined hotspots [39]. Similar limitations were observed in the *cII* dataset, where information on non-hotspot sites was limited [38]. Given these constraints in our non-hotspot data, we refrain from speculating whether this decline in performance solely stems from sequence context or if it’s compounded by the inherent dataset imbalance present in *cII* and *lacZ* datasets.

A recurrent and pivotal parameter during our analysis was tip, denoting the angular displacement of the base pair around its y-axis [52]. Notably, this particular parameter exhibited remarkable prominence, emerging as the preeminent feature within each of the five helical datasets. Alternative rotational parameters, including buckle, opening, and propel also garnered high scores through our combined feature scoring approach. This outcome aligns with our expectations, considering that the collective variation in individual base/base step parameters is likely to exert a substantial impact on overall axis displacement. Our findings resonate with prior molecular modelling studies, which highlight the effects of DNA bending at established sites of DNA damage. Using multivariate statistical analysis, Menzies *et al*. report that buckle and opening parameters significantly contribute to the structural differentiation of adducted sequences in *TP53* [36]. Paul *et al*., using crystallographic and fluorescence lifetime (FLT)-based conformational studies, show that elevated values for slide, roll, and twist parameters can aid DNA opening events during NER, highlighting a distinct interplay between rotational and transitional parameters [35]. As such, we refrain from speculating that rotational parameters alone govern NER functionality, instead highlighting the importance of parameters, including tip, as a potent discriminator of hotspot and non-hotspot sites.

A global analysis of feature importance scores also offered insights into the specific positions of base pairs with significant predictive utility. Results highlighted that bases of intrinsic importance to the classification process were distributed across the entirety of the 23-mer region under analysis. Moreover, each specific gene context exhibited a distinct importance profile, emphasising the context-dependent nature of these critical bases. This observation aligns with previous studies indicating that sequence context, even more distal to adjacent bases, can exert influence over local distortion and, consequently, NER capacity [32,33]. The specific mechanism and extent to which distal sequence affect NER, however, remains a topic of extensive debate.

In the NER pathway, the crucial processes of lesion recognition and initiation are instigated by the heterodimeric XPC–RAD23B–CETN2 complex, hereafter referred to as XPC. Using thermal energy alone, XPC scans the genomic landscape for irregularities by unwinding short segments of DNA in a non-specific manner, a process aptly referred to as the interrogation mode [53]. Local distortion or destabilisation induced by a DNA lesion increases XPC residency at the damage site, allowing for the insertion of functional components and subsequent separation of the helical backbone [54-56]. Lesion recognition is an indispensable, rate-limiting step of NER, and stalled XPC is strictly required for the recruitment of transcription factor, TFIIH [57-59]. Therefore, it is reasonable to consider that various base contexts may impart distinct preferences for strand separation, thus potentially influencing the overall effectiveness of NER and, subsequently, mutational frequency. Paul *et al*., in evaluating the role of sequence context on DNA ‘opening’ by Rad4/XPC, highlight that regions of elevated van der Waals stacking energy may confer resistance to DNA opening [35]. Moreover, it is widely reported that variation in stacking stabilisation among both adduct types and sequence contexts can compensate for destabilising distortions caused by these lesions and in turn, resist NER [60]. Interestingly, it has been found that that duplex stability depends linearly on G/C content, with the free energy of G/C base steps consistently lower than those of A/T or “mixed” base pair steps (with one G-C and one A-T pair) [61]. Given that mutations are AT-biased in eukaryotes, it’s conceivable that regions with high GC content might confer resistance to NER processes due to their enhanced thermodynamic stability, primarily governed by base stacking van der Waals interactions [62-64].

Analysis of *TP53, cII*, and *lacZ* mutational signatures appear to echo these insights. By comparing a 10bp region spanning the lesion site (base pairs 2-12), we observe significant statistical disparities in the G/C content between hotspot and non-hotspot sites in *TP53* and *lacZ* (P < 0.005), (supplementary tables 1,3). In contrast, no such distinction is evident within the *cII* gene (supplementary table 2) potentially contributing to the observed limitations in classifier performance during *cII* deployment. While it is vital to explore the stacking profiles of adduct-bound duplexes in the specific sequence contexts discussed, our results strongly indicate that, in the case of BPDE adducts, there exists a substantial preservation of features distinguishing hotspot and non-hotspot sites, primarily governed by variations in G/C content.

In conclusion, our investigation firmly establishes the efficacy of employing a ML-based approach to not only classify, but also predict biological features from a selectively curated set of helical data. This underscores the practical utility of such methodologies in the analysis of results derived from MD simulations. Within the constraints of the models developed, our findings reveal the significant role of regional helical topology in distinguishing mutation hotspot from non-hotspot sites. Particularly, we showcase that the RF classification of BPDE-bound duplexes is primarily steered by variations in base pair tip, likely stemming from differing thermodynamic stability linked to regional GC content. It’s worth noting that while our analysis has provided insights into the distinctive features influencing mutational patterns in well-studied genes, a broader dataset encompassing a larger variety of sequences is essential to comprehensively understand the precise origins of these mutations. We highlight that this approach is not limited to the genes discussed above and could be applied to any gene subset of interest, including those associated with alternative human diseases. Furthermore, our methodology holds potential for screening mutation hotspot sites to assess the mutational impact of both novel and existing drug compounds. This extends the applicability of our approach beyond fundamental research, making it a valuable tool in drug development and toxicology studies.

## Methods

### DNA Sequences

*TP53, cII*, and *lacZ* were selected as suitable models to explore the influence of sequence context on codon mutability. For each gene context, MD simulations were performed on 25-mer duplex sequences encompassing either mutation hotspot or non-hotspot sites in the presence of the 10S (+)-trans-N2-BPDE-dG adduct (Supplementary Table S1-4). Each sequence consisted of a methylated cytosine at position six, and guanine at position 7. To replicate the homomethylation observed in *TP53*-associated lung cancer, unique structures were generated wherein methylation groups were added to all CpG dinucleotides on both forward and reverse strands. For all sequences, position 7 defines a known location of BPDE binding, and subsequently, the mutation site for any hotspot sequence. Simulations were performed on both adducted and non-adducted duplexes. Sequences were defined as having a mutation hotspot at position 7 if the frequency of G:C>T:A substitutions observed at this site was significantly elevated compared to the expected frequency. *TP53* codon 282 is characterised by a G:C>A:T substitution, however, given its extremely high incidence in lung tumours, was included in our analysis. Mutational data for *TP53, cII*, and *lacZ* gene contexts were obtained from [37], [38], and [39], respectively.

### Starting Structures

All sequences were assembled using the Builder toolbox in PyMOL [65]. This process involved the manual input of each sequence in the form of the single-letter code. Once inputted, Builder then generates the corresponding nucleotide structure, assembling coordinates for sugar, phosphate, and base groups, as well as those for any modifications present. Builder was also utilised in the assembly of the methylated cytosine bases. A nuclear magnetic resonance (NMR) structure for the 7S,8R,dihydroxy-9R,10S-epoxy-7S,8R,9R,10S (+)-trans-anti-B[*a*]PDE DNA adduct (PDB ID: 1AXO) was utilised to assemble carcinogen-bound duplexes [66]. This PDB file contains coordinates for six structures obtained from a relaxation matrix refinement, and the first structure was chosen. The sequence present within the obtained structure was remodelled to resemble hotspot and non-hotspot sites within each gene context using the Builder toolbox as described above.

### Molecular Dynamics Simulations (MD)

All simulations were carried out in triplicate using the GROMACS package and Amber99 force field with backbone torsion modifications, bsc1 [67,68]. Forcefield parameters for BPDE-bound to guanine are detailed in [46]. Partial charges for the methylated cytosine base were calculated using Atomic Charge Calculator and subsequently added to the forcefield [69]. All structures were placed in a cubic box, solvated using the explicit water model TIP3P, and neutralised with the appropriate number of Na^+^□ions prior to simulation. Temperature coupling thermostat was applied using v-rescale, and Particle-mesh Ewald (PME) was used for long-range electrostatics. Simulations were carried out using the NPT ensemble, with periodic boundary conditions, at a temperature of 310.15°K, and a pressure of 1atm. All simulations were performed, in triplicate, using three-stage process: steepest descent energy minimisation with a tolerance of 1000 kJ mol^-1^ nm^-1^, followed by a two-stage equilibration process, each one 50,000 steps in length with a time step integration of 0.002ps, making a total of 100ps; and an MD stage run for a total of 100ns.

### DNA Structural Parameters

To evaluate structural distortion resulting from adduct formation, we assessed seventeen distinct DNA structural parameters for each Molecular Dynamics (MD) trajectory. These parameters encompassed both intra-base pair characteristics—shear, propeller, buckle, stretch, stagger, and opening—as well as inter-base pair attributes—twist, roll, tilt, rise, shift, and slide. Supplementing these, two translational base pair axis parameters (X and Y displacement), two rotational parameters (inclination and tip), and an axis-bend parameter were calculated to define base pair geometry relative to the helical axis. Parameters were measured using open source Curves+ and Canal□software [52].

### Classifier Comparison

To select an optimal classifier, we compared the predictive capabilities of 15 distinct classification algorithms. We compared the performance of Nearest Neighbours, Linear support vector machine (SVM), Polynomial SVM, Radial Basis Function (RBF) SVM, Gaussian Process, Gradient Boosting, Decision Tree, Extra Trees, Random Forest, Neural Network, AdaBoost, naïve Bayes, Quadratic Discriminant Analysis (QDA), Stochastic Gradient Descent (SGD), and Linear Discriminant Analysis (LDA). Each classifier was instantiated with default parameters and trained on 80% of the *TP53* dataset. Trained models were then used to predict the target variable for the testing set, with metrics accuracy, precision, recall, f1, train-test gap, and AUC ROC used to evaluate model performance. Helical information for base pairs 3-23 only was used as ML input data.

### Random Forest

Each decision tree in the RF was trained to predict the repair phenotype and carcinogen status of the duplexes described above, based on their structural dynamics (Figure 5). For model evaluation, 80% of the *TP53* dataset was randomly selected as training data, with the remaining data being separated for model testing. Utilising identical training events, the classifier was then deployed to various mutational models including *cII, LacZ*, and novel *TP53* datasets, where 95% of the unseen data was used for testing, ensuring a comprehensive representation of RF applicability. Optimal hyperparameters were selected using 10-fold GSO. Each RF model were implemented using the RandomForestClassifier module from the Python machine learning library scikit-learn [70]. RF models were ran using the same 10 random seeds to ensure reproducibility across each validated model.

**Figure 5:**
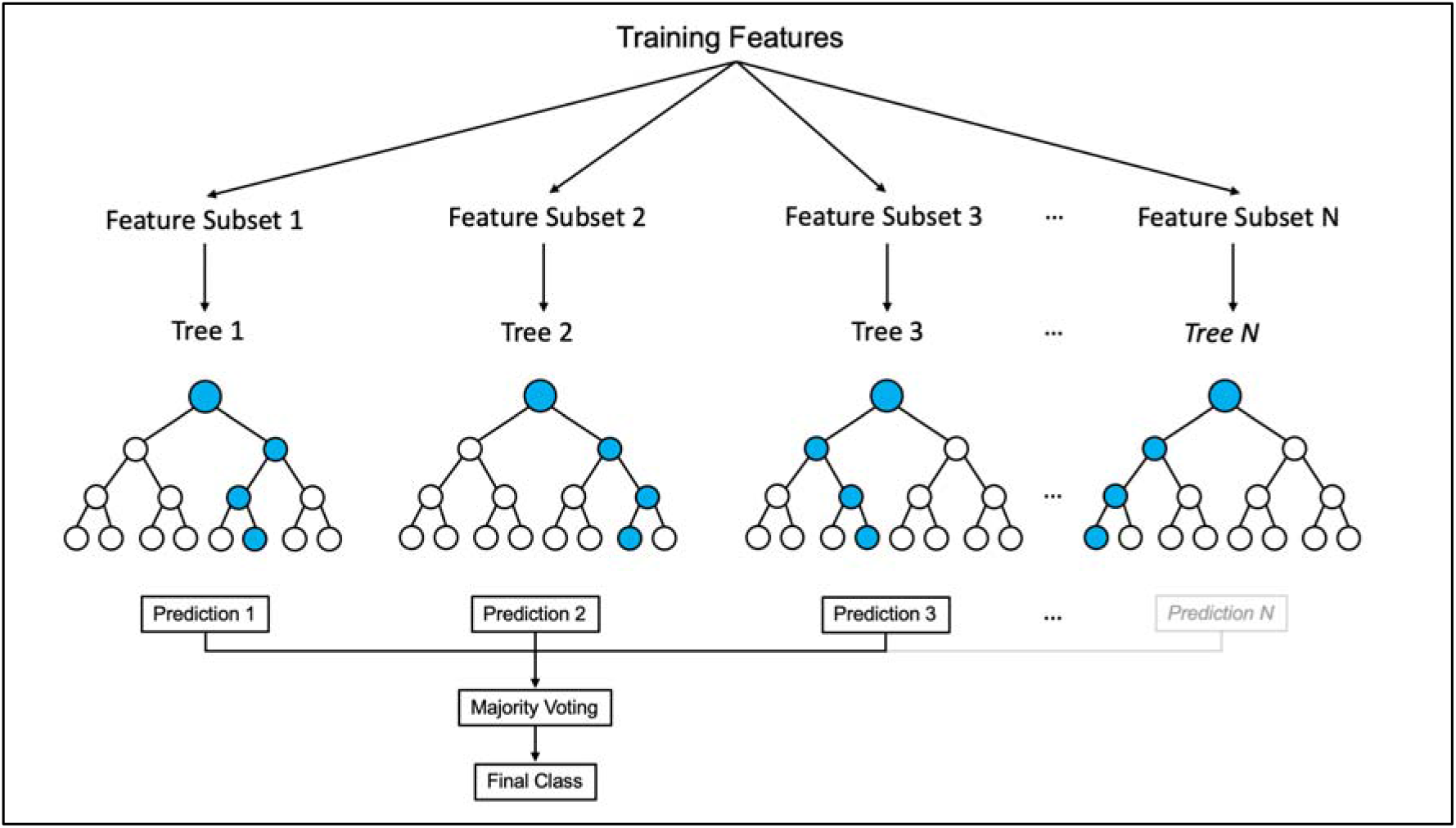
Framework of a Random Forest algorithm.

Feature selection was conducted to identify a subset of relevant features essential for constructing robust learning models and informing the classification algorithm. Multiple feature selection techniques were employed to ensure comprehensive coverage and insights into feature importance. Leveraging RF feature importance, recursive feature elimination, and lasso regularization, we identified the most informative features for our analysis. From each technique, we extracted the top 10% (36) features. These features were assigned scores ranging from 36 (highest importance) to 0 (lowest importance), excluding features with a score of 0. The resultant scores from each technique were amalgamated across datasets, resulting in a comprehensive list of the most significant features for our investigation.

