## Supplementary figures for "Machine learning to classify mutational hotspots from molecular dynamic simulations"

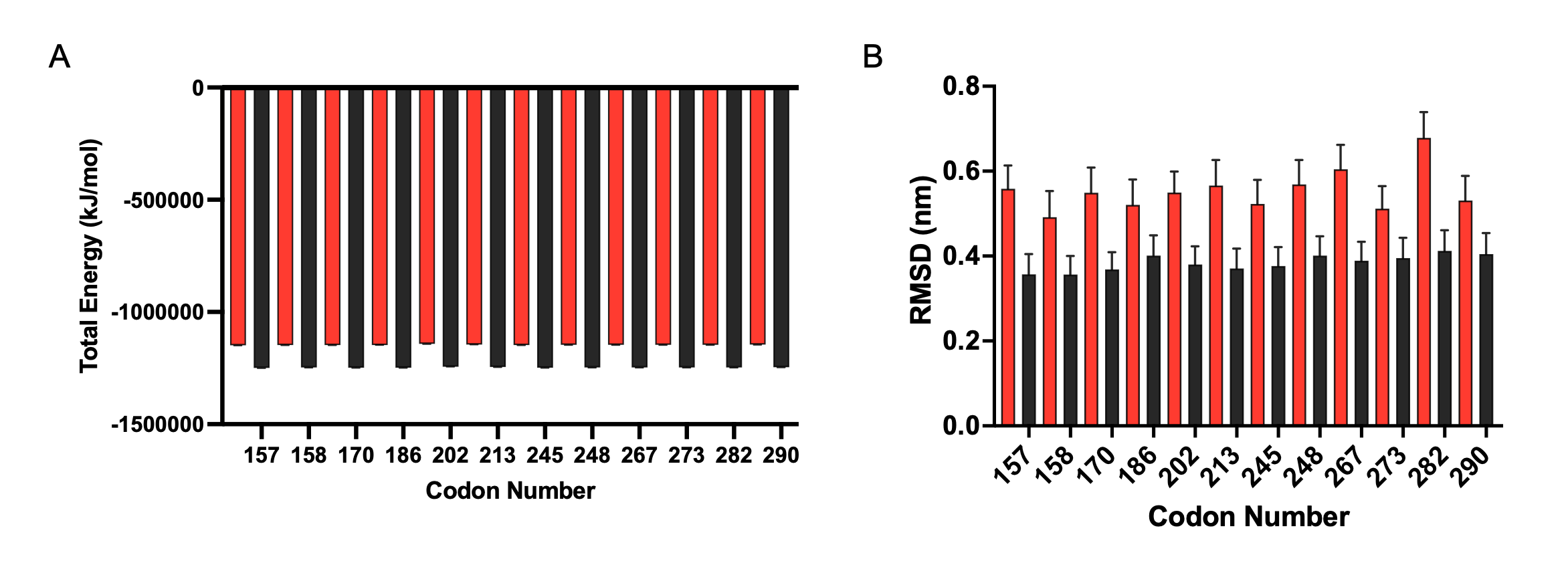

*Supplementary Figures S1: Conformational stability and flexibility of adducted and control Val duplexes. Total energy (****A****) in KJ/mol and RMSD values (****B****) in nm for control sequences (black) and adducted sequences (red); error bars show ± standard deviation.*

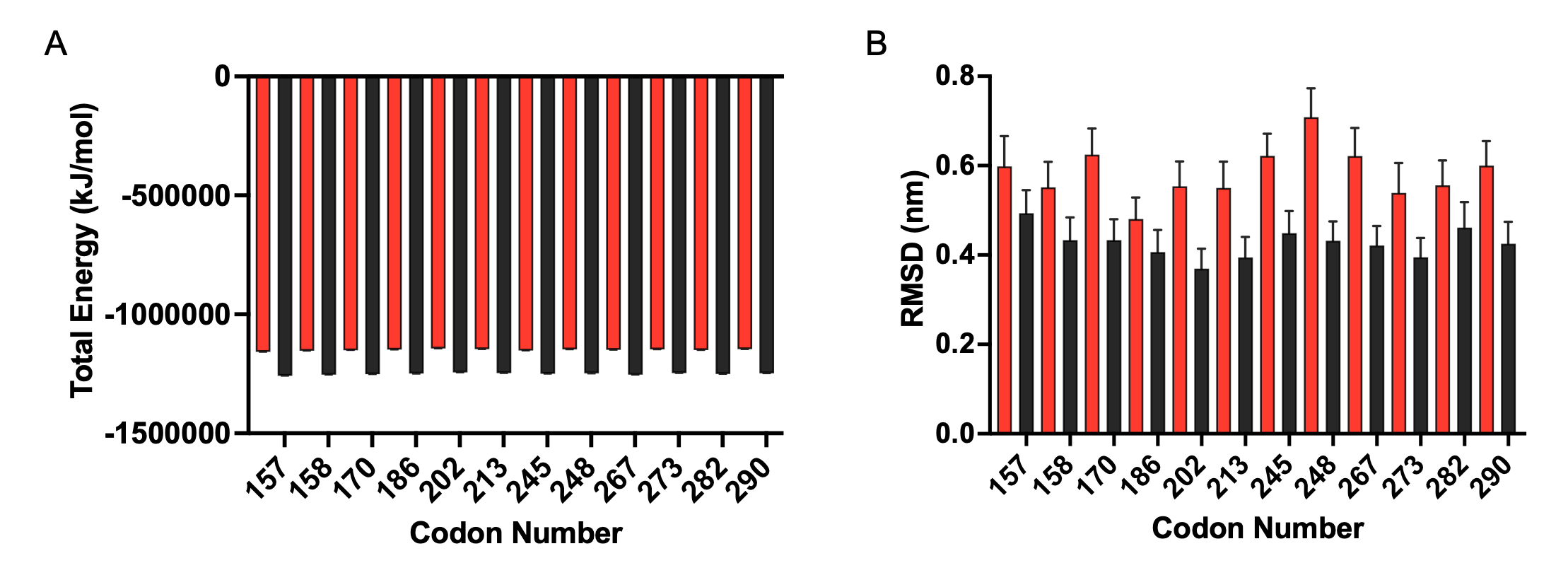

*Supplementary Figures S2: Conformational stability and flexibility of adducted and control Meth duplexes. Total energy (****A****) in KJ/mol and RMSD values (****B****) in nm for control sequences (black) and adducted sequences (red); error bars show ± standard deviation.*

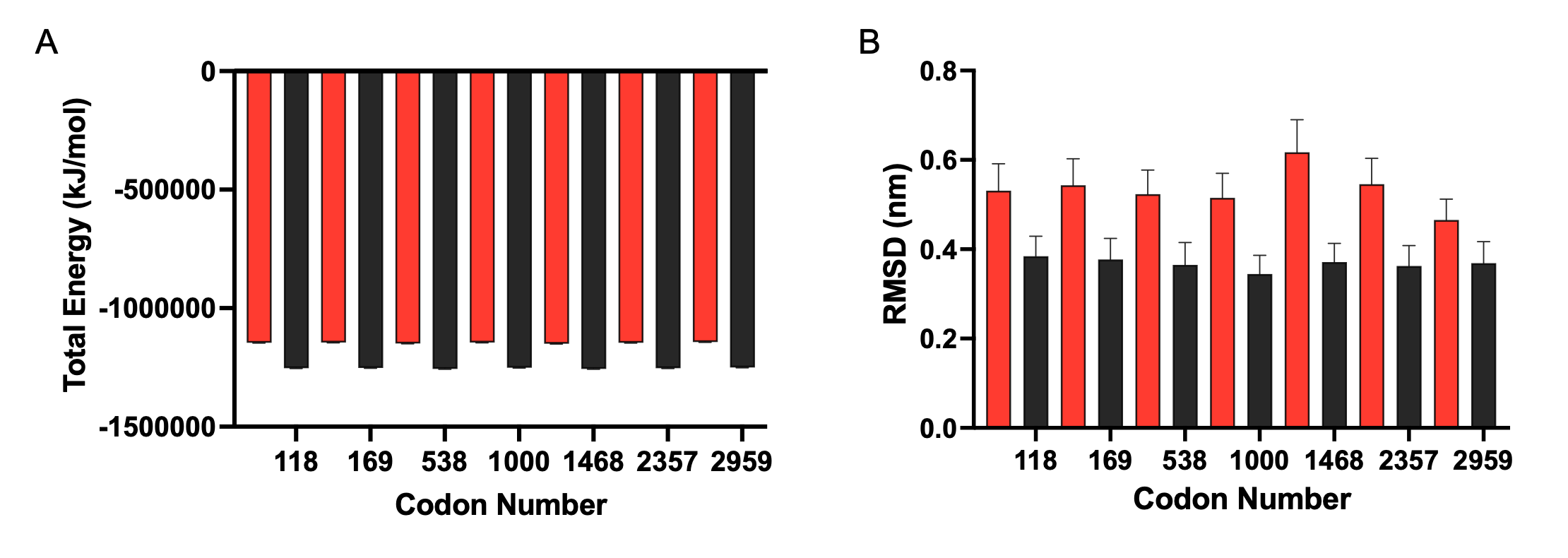

*Supplementary Figures S3: Conformational stability and flexibility of adducted and control lacZ duplexes. Total energy (****A****) in KJ/mol and RMSD values (****B****) in nm for control sequences (black) and adducted sequences (red); error bars show ± standard deviation.*

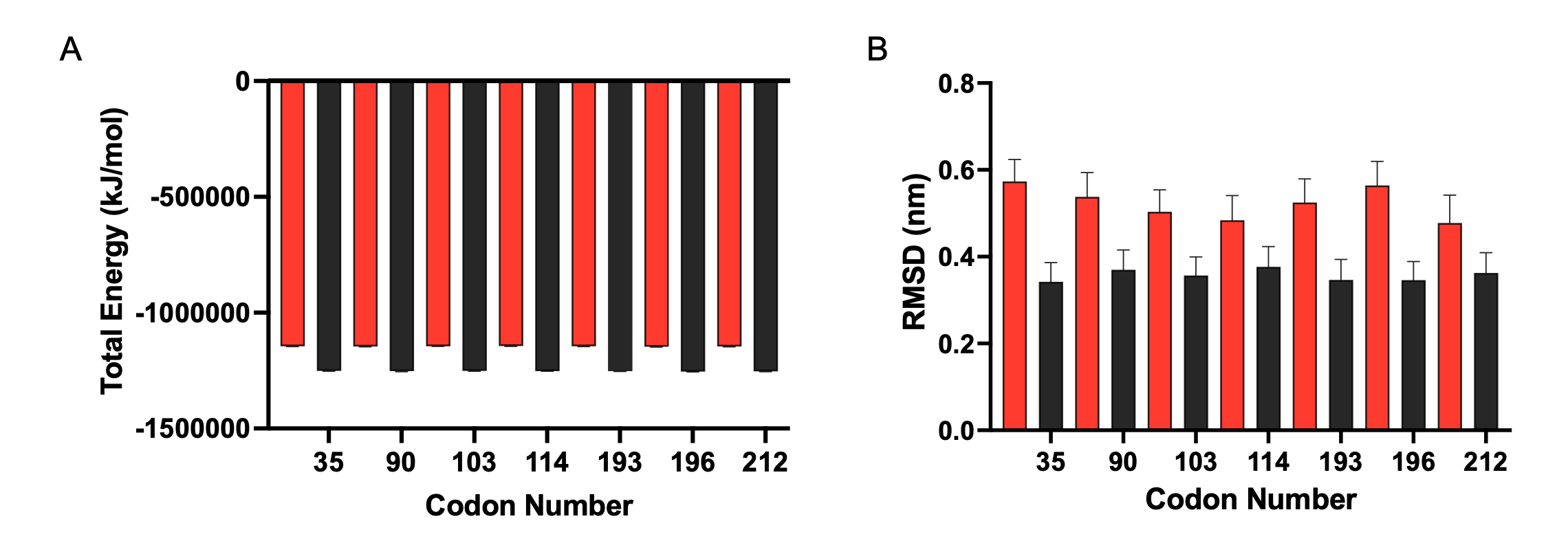

*Supplementary Figures S4: Conformational stability and flexibility of adducted and control cII duplexes. Total energy (****A****) in KJ/mol and RMSD values (****B****) in nm for control sequences (black) and adducted sequences (red); error bars show +/− standard deviation.*

*Supplementary* *Table 1:* TP53 *DNA sequences used for molecular dynamics simulations (adducted guanine underlined). Sequences were determined to be hotspots in cancers based on their prevalence in the COSMIC database. Difference between GC content of hotspot and non-hotspot sites (base pairs 2-12) was statistically significant (P < 0.005).*

| Codon | Sequence | Hotspot Status (across cancers) | No. observed G:C>T:A in lung cancer | GC Content (%) |
| --- | --- | --- | --- | --- |
| 157 | ACCCGC*GTCCGCGCCATGGCCATCT | Lung | 120 | 72.0 |
| 158 | GCGTCC*GCGCCATGGCCATCTACA | Lung | 107 | 66.7 |
| 245 | ATGGGC*GGCATGAACCGGAGGCCCA | Many | 108 | 68.0 |
| 248 | TGAACC*GGAGGCCCATCCTCACCAT | Many | 75 | 60.0 |
| 273 | AGGTGC*GTGTTTGTGCCTGTCCTGG | Many | 134 | 60.0 |
| 282* | GAGACC*GGCGCACAGAGGAAGAGAA | Many | 1 | 60.0 |
| 170 | CATGAC*GGAGGTTGTGAGGCGCTGC | None | 0 | 64.0 |
| 186 | GATAGC*GATGGTCTGGCCCCTCCTC | None | 0 | 64.0 |
| 202 | ATTTGC*GTGTGGAGTATTTGGATGA | None | 0 | 40.0 |
| 213 | CTTTTC*GACATAGTGTGGTGGTGCC | Breast | 11 | 52.0 |
| 267 | TGGGAC*GGAACAGCTTTGAGGTGCG | None | 2 | 60.0 |
| 290 | ATCTCC*GCAAGAAAGGGGAGCCTCA | None | 0 | 56.0 |

*Sequences displaying majority substitutions distinct from G:C>T:A.

*Supplementary* *Table 2:* cII *DNA sequences used for molecular dynamics simulations (adducted guanine underlined). Sequences were determined to be hotspots in cancers based on the mutational frequency of the* cII *transgene in embryonic mouse fibroblasts upon exposure to BPDE. Difference between GC content of hotspot and non-hotspot sites (base pairs 2-12) was not statistically significant (P > 0.05).*

| Codon | Sequence | Hotspot Status | GC Content (%) |
| --- | --- | --- | --- |
| 35 | CTCTAC*GAATCGAGAGTGCGTTGCT | Hotspot | 52.0 |
| 103 | GTGGGC*GTTGATAAGTCGCAGATCA | Hotspot | 52.0 |
| 193 | GTTGAC*GACGACATGGCTCGATTGG | Hotspot | 56.0 |
| 196 | GACGAC*GACATGGCTCGATTGGCGC | Hotspot | 64.0 |
| 212 | TGTCGC*GCCAATCGAGCCATGTCGT | Hotspot | 60.0 |
| 90 | GACAGC*GGAAGCTGTGGGCGTTGAT | Non-Hotspot | 60.0 |
| 114 | TAAGTC*GCAGATCAGCAGGTGGAAG | Non-Hotspot | 52.0 |

*Supplementary* Table 3: lacZ *DNA sequences used for molecular dynamics simulations (adducted guanine underlined). Sequences were determined to be hotspots in cancers based on their mutational frequency by high throughput next generation sequencing. Difference between GC content of hotspot and non-hotspot sites (base pairs 2-12) was statistically significant (P < 0.005).*

| Codon | Sequence | Hotspot Status | No. observed G:C>T:A upon BPDE exposure | GC Content (%) |
| --- | --- | --- | --- | --- |
| 169 | AATGGC*GAATGGCGCTTTGCCTGGT | Hotspot | 21 | 56.0 |
| 538 | CGCGCC*GGAGAAAACCGCCTCGCGG | Hotspot | 16 | 76.0 |
| 1000 | TTCCGC*GAGGTGCGGATTGAAAATG | Hotspot | 36 | 52.0 |
| 2468 | GGCGGC*GGAGCCGACACCACGGCCA | Hotspot | 17 | 80.0 |
| 2357 | TCACCC*GTGCACCGCTGGATAACGA | Hotspot | 26 | 60.0 |
| 118 | AATAGC*GAAGAGGCCCGCACCGATC | Non-Hotspot | 3 | 60.0 |
| 2959 | AATATC*GACGGTTTCCATATGGGGA | Non-Hotspot | 3 | 44.0 |

*Supplementary Table 4: Helical parameters and their abbreviations.*

| Parameter | Abbreviation | Description |
| --- | --- | --- |
| Base Pair Step Parameters | | |
| Twist/Rise | tw/re | Rotation and Translation of the base pairs about an axis which is perpendicular to the plan of the base step |
| Roll/Slide | rl/se | Rotation and Translation of the base pair about an axis which is the long axis of the base step |
| Tilt/Shift | tt/st | Rotation and Translation of the base pair about an axis which is the short axis of the base step |
| Base Pair Parameters | | |
| Opening/Stagger | og/sg | Rotation and Translation of the bases about an axis which is perpendicular to the plane of the base pair |
| Propeller/Stretch | pl/sh | Rotation and Translation of the bases about an axis which is the long axis of the base pair |
| Buckle/Shear | be/sr | Rotation and Translation of the bases about an axis which is the short axis of the base pair |
| Axis-Displacement Parameters | | |
| X and Y Displacement | xp/yp | Displacement of the bases in the x or y direction |
| Axis-bend | ad | Distance measure of the axis-bend |
| Incline/Tip | in/tp | Angle movement across the x and y axis |
